# Stimulus-specific recruitment of human amygdala neurons predicts episodic memory encoding success

**DOI:** 10.1101/2025.10.29.685162

**Authors:** Jiayang Xiao, Jonathan Daume, Yousef Salimpour, Natalia Kurilenko, Clayton Mosher, William S. Anderson, Taufik A. Valiante, Adam N. Mamelak, Ueli Rutishauser

## Abstract

Controlling whether a given experience is encoded into long-term memory and thus later remembered is a crucial component of our memory system whose failure is often at the root of memory disorders. One brain area that takes part in controlling which experiences are remembered is the amygdala, but the mechanisms by which it does so remain poorly understood. Here we examined single-neuron activity and local field potentials as human participants performed recognition memory tasks with visual stimuli. Category-selective amygdala neurons exhibited elevated firing rates during encoding of later remembered items versus forgotten items. This subsequent memory effect was restricted to images of the preferred category of a given cell, was stronger and appeared earlier in the amygdala compared to the hippocampus, and did not depend on the valence and arousal of the stimuli. In contrast, category selective cells immediately upstream in the ventral temporal cortex did not exhibit a subsequent memory effect, highlighting specificity to the amygdala. Successful memory formation was accompanied by enhanced spike-field coherence between the activity of category cells in the amygdala and hippocampal field potentials. These findings, replicated in two large independent datasets with two different tasks, demonstrate that recruitment of stimulus-specific amygdala representations predicts episodic memory formation, particularly in the right amygdala. This data suggests category cells in the right amygdala as a cellular target for interventions to treat memory disorders in humans.

## Introduction

In a typical day, we experience a large number of events, each accompanied by large amounts of sensory information. Imagine walking through a farmers’ market: the sounds of people talking, the faces in the crowd, the scent of freshly cooked food from nearby stalls, and the taste of an exceptionally delicious dish, all stimulate your senses. Despite this rich sensory experience, typically only a small and select subset of this experience is transformed into long-lasting episodic memories. The neural mechanisms that determine whether a long-term memory of an experience is created, and if so what aspects of the experience are encoded, are only beginning to be understood^1–4^. A major hypothesis is that only some experiences are ‘tagged’ for consolidation into long-term memory^5–7^, with the remainder only residing briefly in short-term memory. A brain area that plays a major role in memory tagging is the amygdala, which is thought to increase memory encoding and/or consolidation for emotionally significant events by modulating synaptic plasticity processes taking place in the hippocampus^8–13^. While classically thought to be restricted to highly emotional stimuli, modifying overall amygdala activity around the time of experiencing novel sensory input enhances the memory of a wide range of neutral stimuli despite not evoking a physiological emotional response^14–17^. While these findings establish the amygdala as a powerful modulator of memory encoding (and thus establishing it as a potential target for a memory prosthesis), little is known about the underlying mechanisms. In particular, it remains unclear whether the amygdala is involved in processing the content of the to-be-encoded memory itself or whether its role is to modulate attention or arousal, without processing the content of the memory itself. Here, we utilize large-scale single-neuron recordings in the human brain in two different memory tasks to examine this important open question.

A key motivation for identifying mechanisms that control memory encoding processes is that modulating such mechanisms could be a way to treat memory disorders. In this context, the amygdala has emerged as a promising therapeutic target, with several studies showing ability to enhance memory encoding without inducing emotional responses by stimulating the amygdala^14,15,17^. This is in stark contrast to hippocampal stimulation, which in many circumstances disrupts rather than enhances memory formation^18–21^. While the amygdala is a promising new target for a memory prosthesis, the lack of an underlying mechanism leaves it unclear what exactly the therapeutic target is in the amygdala for enhancing memory. Is it to increase or decrease the overall activity of all cells regardless of their location and tuning properties, or is it rather to modify the activity of specific cells while leaving others unchanged? Here, we address this major open question by comparing the response of cells in the human amygdala to novel stimuli between those that were later remembered and those that were forgotten (difference due to memory approach^22–25^). The amygdala receives prominent inputs from several high-level sensory areas^26,27^ and contains well-characterized high-level representations of memory content^28–30^. Based on this body of literature, here we hypothesize that what predicts memory encoding success is the selective recruitment of amygdala cells that represent aspects of the to-be-encoded memory content rather than overall changes of activity. We tested this overarching hypothesis in two separate and distinct datasets (“Task 1” and “Task 2”), allowing us to assess the robustness of the effect. If supported, this would mean that for a therapeutic intervention to be effective, it would have to increase the gain of existing stimulus-specific responses rather than to change overall levels of activity.

## Results

### Behavior (Task 1)

We recorded single-neuron activity in neurosurgical patients while they performed a working memory task followed by a recognition memory test (42 patients, 49 sessions, Fig. 1a). In each trial of the working memory task (Fig. 1b), patients viewed one or three pictures and, after a maintenance period, were asked whether the probe picture matched (was identical) to one of the 1–3 images seen during encoding in that trial. New, never-before-seen pictures were shown during encoding in each of the 140 trials, allowing us to later examine long-term memory for the same images. The probe image was always an image seen before, either in the current trial or in a previous trial. Each image belonged to one of five visual categories (people, food, animals, cars, and landscapes). After a 10–30 minute long delay period following the completion of the working memory task, patients performed a long-term memory test to assess whether they recognized the images they saw earlier during the working memory part of the task (Fig. 1c). Patients were shown 400 images, 200 of which were shown before during the working memory part and thus familiar, and 200 of which were novel (never shown so far). For each image, patients were asked to indicate whether they had seen the picture before (‘new’ or ‘old’) and how sure they were of their decision (Fig. 1c).

**Fig. 1:**
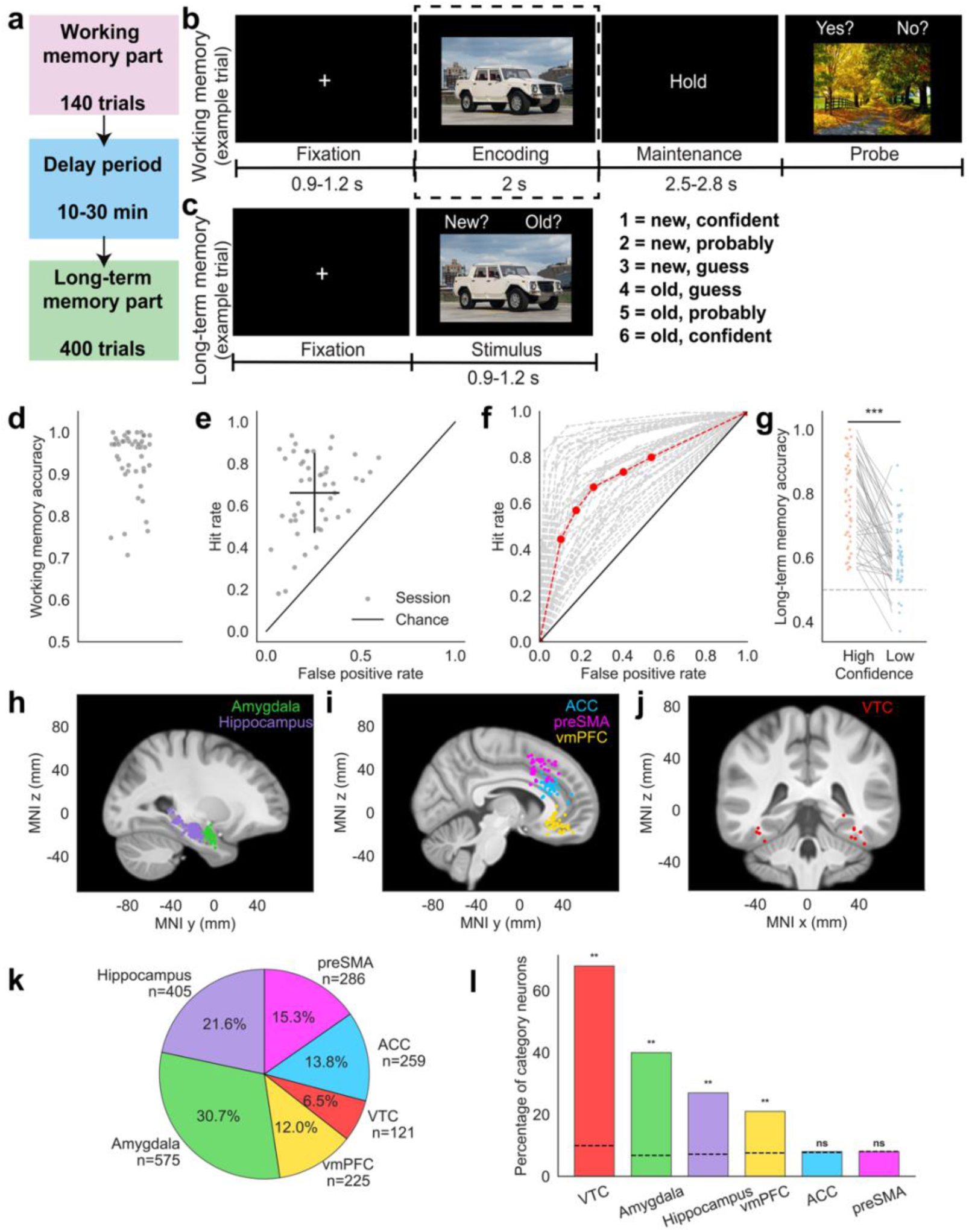
Task, behavior, and recording sites (Task 1). **(a)** Task. This experiment consisted of two parts: a working memory part and a long-term memory part, separated by a 10–30 minute delay. **(b)** In the working memory part, participants viewed either one or three novel, never-before-seen pictures and after a maintenance period, indicated whether a probe picture was previously seen on the same trial or not. All pictures belonged to five different categories. **(c)** The long-term memory part required participants to assess whether each picture presented was ‘new’ (never seen before) or ‘old’ (previously seen in the working memory part) together with a confidence rating. **(d-g)** Behavior. **(d)** Accuracy of responses in the working memory shows subjects had good working memory (each dot is a session, chance is 50%). **(e-g)** Behavior in the long-term memory part shows subjects had good long-term memory. **(e)** Hit rate and false positive rate. Each point is one session, black lines are the mean performance ± SD. **(f)** Behavioral receiver operating characteristic (ROC) curves for individual sessions (grey) and the average (red). Each data point along the curve corresponds to a specific confidence level, beginning with the highest (6: old, confident) located in the lower-left corner. The diagonal line indicates chance level. The performance was above chance at all levels of confidence, with a mean area under the curve of 0.73 ± 0.11 (mean ± SD). **(g)** Recognition memory performance was significantly higher in high vs. low-confidence trials. Each line connects the two dots belonging to the same session (Chance level = 50%, Wilcoxon signed-rank test, *** indicates *p* < 0.001). **(h-j)** Electrode locations. Each dot denotes a microwire bundle. Coordinates are in Montreal Neurological Institute (MNI) space. Left and right hemispheres are collapsed in panels h and i. **(k)** Distribution of recorded neurons. **(l)** Proportion of cells within each brain region that qualified as category cells. Bars represent observed values, and horizontal lines indicate the 99th percentile of the surrogate null distribution. Statistical significance is denoted as follows: *p* < 0.01 (**), not significant (ns).

Patients had good working memory, with average accuracy of 93.4% ± 7.5% (mean ± SD, chance level = 50%, *p* < 0.001, Wilcoxon signed-rank test) for answering the probe question (Fig. 1d). Patients also had good long-term memory: their accuracy in the later recognition memory test was 70.3% ± 9.7% (Fig. 1e, chance level = 50%, *p* < 0.001, Wilcoxon signed-rank test). Furthermore, accuracy on the recognition memory test was correlated with confidence, with higher confidence ratings corresponding to more accurate decisions (Fig. 1f-g; see legend for statistics). This indicates that subjects formed declarative memories for the stimuli^31^.

### Category-selective neurons in the amygdala predicts memory encoding success (Task 1)

Neural recordings in Task 1 yielded 1871 well-isolated single neurons, distributed across the amygdala (575 neurons), hippocampus (405 neurons), anterior cingulate cortex (ACC, 259 neurons), presupplementary motor area (preSMA, 286 neurons), ventromedial prefrontal cortex (vmPFC, 225 neurons) and ventral temporal cortex (VTC, 121 neurons) (Fig. 1h-k).

We first identified neurons whose response following stimulus onset varied significantly across the five different visual categories from which all images were chosen. To identify these “category” neurons, we conducted a 1 × 5 ANOVA on the average firing rate within a 200 to 1000 msec window after stimulus onset. We included spike counts pooled across all encoding periods and the probe period of the working memory task. We then performed a right-sided permutation-based t test to compare the category with the highest average firing rate to all other categories. A neuron was considered a category neuron if both tests were significant with *p* < 0.05, with the category showing the highest average firing rate being identified as the preferred category for that neuron. To determine the chance level, the same selection process was repeated 1000 times using randomly shuffled category labels, generating a null distribution of category neuron proportions within each region. The proportion of category neurons out of all recorded neurons in a given brain area was higher than expected by chance in the VTC (67.8%), amygdala (40.0%), hippocampus (27.2%), and vmPFC (21.3%), but not in the ACC or pre-SMA (permutation test, *p* < 0.01, Fig. 1l). We focused on the four regions with a significant proportion of category neurons for the analysis that follows.

As expected, category neurons showed an increase in firing rate following the onset of an image that belonged to the preferred category of the cell (Fig. 2a-b shows two examples, left side). We hypothesized that if this increase in firing rate is related to memory formation, its extent should be predictive of memory strength as assessed using the later recognition memory test. To test this hypothesis, we categorized all instances during which an image was seen for the first time (when it is novel) as ‘remembered’ or ‘forgotten’ based on whether the behavioral response was ‘old’ or ‘new’ to the same picture (when it is familiar) during the later recognition memory test. Comparing trials of the preferred category between these two types revealed that this was indeed the case for the two example neurons shown (Fig. 2a-b, right side). For example, the neuron that increased its firing most to pictures showing animals exhibited stronger firing for images in this category that were later remembered compared to those that were forgotten (Fig. 2a). A similar pattern was observed for the example neuron that preferred the ‘food’ category (Fig. 2b).

**Fig. 2:**
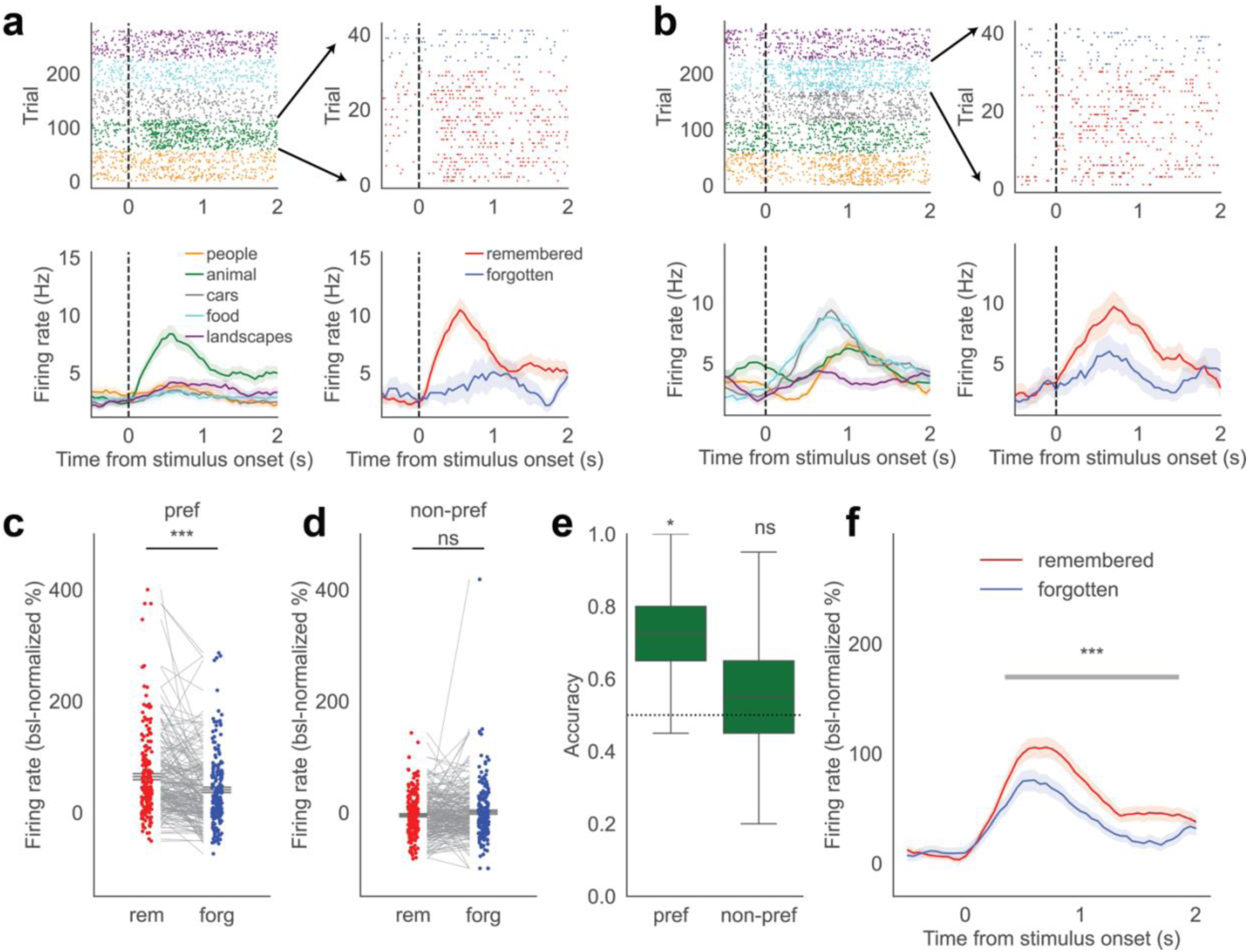
Amygdala activity during encoding in Task 1 differs between later remembered and later forgotten stimuli. **(a-b)** Two example amygdala neurons during memory encoding. For each neuron, the left plot shows category selectivity, with increased firing rates for the preferred image category. The right plot shows that the same neuron exhibited higher firing rates for images from the preferred category that were later remembered compared to those that were forgotten. **(c)** Comparison of the average firing rate during the encoding period between later remembered and forgotten trials in the preferred category in the amygdala. Each line connects the two dots belonging to the same neuron. Horizontal lines indicate mean ± SEM. Statistical analysis was performed using the Wilcoxon signed-rank test. Amygdala: n = 196 neurons, *p* = 4.3 × 10^−8^, *p* value was Bonferroni corrected for all four brain regions with significant proportions of category cells. Neurons were excluded if they had insufficient data in any condition or exhibited activity more than four standard deviations from the average. **(d)** Same as (c), but showing the average firing rate for trials in the non-preferred category. Amygdala: n = 208 neurons, *p* = 0.44, Bonferroni corrected for multiple comparisons. **(e)** Population decoding of remembered versus forgotten items based on activity in the amygdala. Boxplots show the 25th and 75th percentiles, with horizontal lines marking the medians and whiskers extending to 1.5 times the interquartile range. The dotted line represents chance-level accuracy. The left box shows decoding performance for trials from the preferred picture category (*p* = 0.017), and the right box shows performance for the non-preferred category (*p* = 0.25). **(f)** Time-resolved firing rate in the preferred category, averaged across all amygdala category-selective neurons. Horizontal bar depicts time points where the firing rate for the preferred category differs significantly between later remembered and forgotten trials (cluster-based permutation test). Data are presented as mean values ± SEM. \**p* < 0.05, \*\**p* < 0.01, \*\*\**p* < 0.001, ns indicates non-significant.

To quantify this effect at the population level across all identified category cells, we calculated the average firing rates of category neurons from 200 to 2000 msec after stimulus onset, separately for trials belonging to the preferred and non-preferred categories of each cell. This revealed that, during the encoding period, the average firing rates in the amygdala were significantly higher for remembered trials than for forgotten trials, but only for trials from the preferred category (Fig. 2c-d, Wilcoxon signed-rank test, *p* = 4.3 × 10^−8^ for preferred category vs. *p* = 0.44 for non-preferred category in the amygdala after Bonferroni correction for four regions).

We next used a generalized linear mixed-effects model (GLM) to assess the strength of the relationship between recognition memory accuracy and firing rates of category neurons during encoding in the amygdala. Firing rate was the dependent variable, later memory accuracy (remembered vs. forgotten) was the predictor variable, and neuron ID nested within recording session ID were included as random effects. This analysis, which included only preferred trials, revealed a significant effect of memory accuracy (β = 23.73, t = 4.60, *p* = 4.3 × 10^−6^), suggesting that neuronal activity of category cells in the amygdala is predictive of later memory performance. In contrast, no significant effect of memory accuracy on the firing rate of amygdala cells was observed for non-preferred trials (β = −3.31, t = −1.29, *p* = 0.20). Amygdala neurons predicted memory formation also when only considering the first picture in each trial (“encoding 1”), as well as when considering each of the five visual categories separately (Supplementary Fig. 1). Together, this data shows that category cells in the amygdala show a robust subsequent memory effect for the subset of stimuli belonging to their preferred category.

Could this effect also be seen when considering all trials across all recorded amygdala cells? This was the case: there was a significant positive relationship between mean firing rate and later retrieval success or failure across all trials (β = 4.70, t = 3.30, *p* = 9.5 × 10^−4^, n = 496 cells included), indicating that on average, higher firing rates of amygdala cells are associated with a higher likelihood of successfully forming a long-term memory. However, this effect was driven by the preferred-category trials of category cells: when repeating above analysis after removing, for all cells, the trials of the category to which the response was maximal of that cell, no significant effect remained (β = 2.30, t = 1.57, *p* = 0.12, n = 496 cells). This indicates that the overall memory-related modulation was driven primarily by responses to preferred stimuli of category cells, indicating that these cells specifically are part of the mechanism for modulating memory encoding strength.

To test whether long-term memory accuracy can be predicted using neural activity during encoding, we performed population decoding using the firing rates of category-selective neurons in the amygdala during encoding (cross-validated). Consistent with our GLM-based results, we found that based on amygdala activity during encoding, subsequent memory outcomes for the subset of images belonging to a neuron’s preferred, but not the non-preferred category, could be predicted for single trials with accuracy significantly higher than expected by chance (Fig. 2e, mean accuracy = 72.4%, *p* = 0.017 for preferred category, mean accuracy = 56.2%, *p* = 0.25 for non-preferred category; chance is 50%).

At what point of time was neural activity predictive of memory formation? Neurons in the amygdala exhibited higher firing rates for remembered items within their preferred category from 350 to 1850 msec after stimulus onset (Fig. 2f, cluster-based permutation test, *p* < 0.001). Since the average differential response latency of amygdala category cells is approximately 300 msec^31,32^, this finding indicates that the selective recruitment of cells for memory encoding is nearly instantaneous with the sensory response.

### Limited subsequent memory effects outside the amygdala

Did category cells in other brain areas also exhibit subsequent memory effects? Other than the amygdala, significant proportions of cells qualified as category cells in the hippocampus, VTC, and vmPFC, allowing us to answer this question. Similar to cells in the amygdala, hippocampal category cells also fired more during the encoding of items from the preferred category when those items were later remembered compared to when they were not (Fig. 3a, *p* = 0.019 for hippocampus after Bonferroni correction for multiple brain regions). The subsequent memory effect observed in the hippocampus was also specific to the preferred category and was not significant during the encoding of items from the non-preferred category (Fig. 3c, Wilcoxon signed-rank test, *p* > 0.05 for all regions). In contrast, no significant subsequent memory effect was observed for category cells recorded in the VTC or vmPFC. Note that this was the case despite the presence of category neurons in those regions (Fig. 3a, Wilcoxon signed-rank test, *p* = 0.91 for VTC, *p* = 0.57 for vmPFC). Indeed, the by far largest proportion of category cells (67.8%) was observed in the VTC, and these cells responded much more strongly than in all the other areas (Fig. 3e). This finding indicates that category responses in the amygdala, which receives direct input from the VTC^33^, are modulated by memory encoding processes in addition to reflecting sensory input. In contrast, in the VTC, the responses were not modulated by memory encoding processes.

**Fig. 3:**
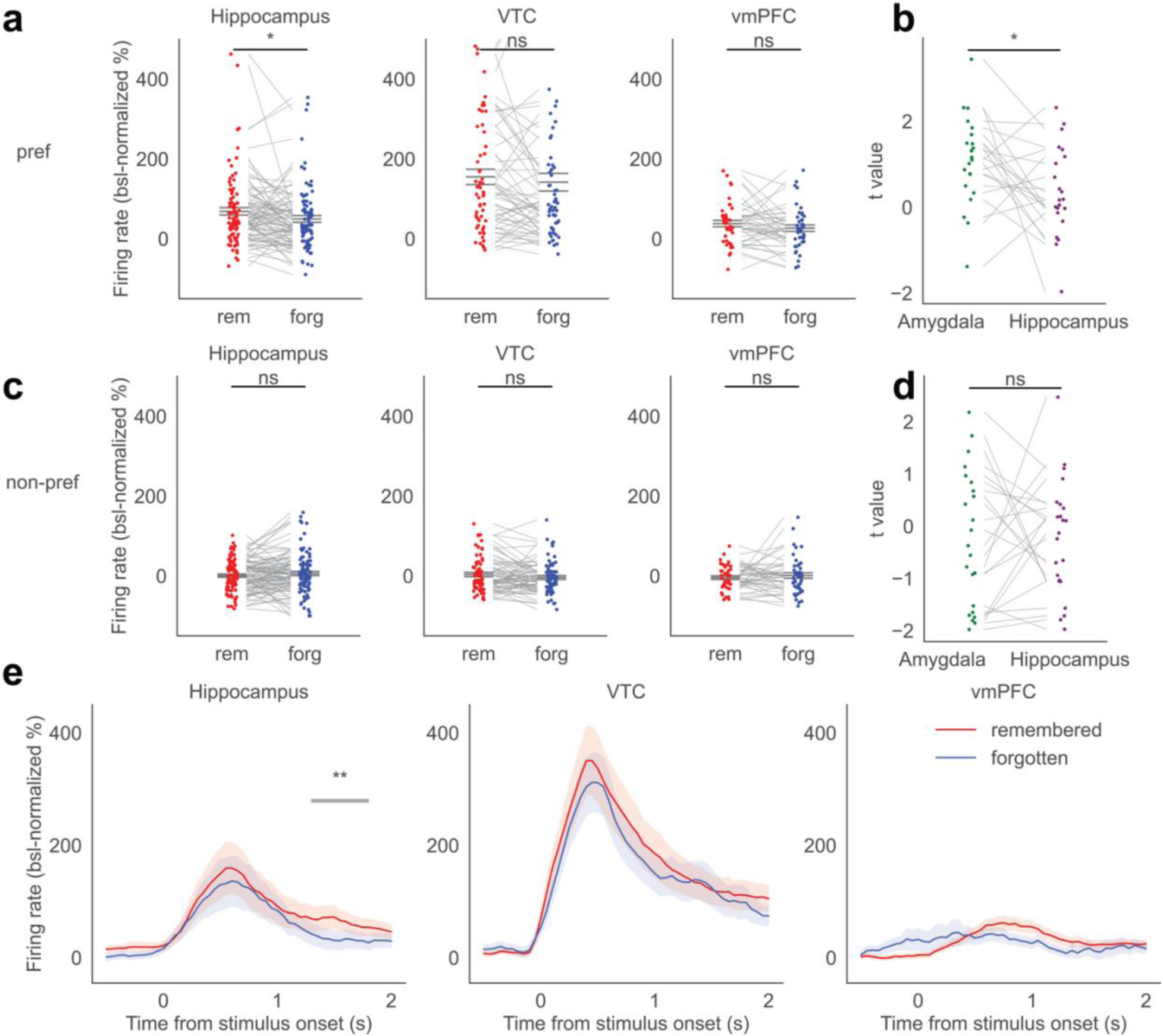
Subsequent memory effect is delayed and weaker in the hippocampus and absent in the VTC and vmPFC. **(a)** Comparison of the average firing rate during the encoding period between later remembered and forgotten trials in the preferred category in the hippocampus, VTC, and vmPFC. Each line connects the two dots belonging to the same neuron. Horizontal lines indicate mean ± SEM. Statistical analysis was performed using the Wilcoxon signed-rank test. Hippocampus: n = 91 neurons, *p* = 0.019; VTC: n = 58 neurons, *p* = 0.91; vmPFC: n = 43 neurons, *p* = 0.57, *p* values were Bonferroni corrected for all four brain regions. Neurons were excluded if they had insufficient data in any condition or exhibited activity more than four standard deviations from the average. **(b)** Comparison of the subsequent memory effect between the amygdala and hippocampus. T-values were obtained by fitting GLM models to data from either the amygdala or hippocampus in each session. Only sessions that had data from both regions were included in this analysis. Each line connects the two data points from the same session. Statistical significance was assessed using the Wilcoxon signed-rank test (n= 22 sessions, *p* = 0.021). **(c)** Same as (a), but showing the average firing rate for trials in the non-preferred category. Hippocampus: n = 97 neurons, *p* = 0.47; VTC: n = 68 neurons, *p* = 0.37; vmPFC: n = 45 neurons, *p* > 0.9, Bonferroni corrected for multiple comparisons. **(d)** Same as (b), but there was no regional difference in the non-preferred category. Statistical significance was assessed using the Wilcoxon signed-rank test (n= 23 sessions, *p* > 0.9). **(e)** Time-resolved firing rate in the preferred category, averaged across all category-selective neurons. Horizontal bar depicts time points where the firing rate for the preferred category differs significantly between later remembered and forgotten trials (cluster-based permutation test). Data are presented as mean values ± SEM. \**p* < 0.05, \*\**p* < 0.01, \*\*\**p* < 0.001, ns indicates non-significant.

To compare the magnitude of the memory encoding effect between the amygdala and the hippocampus, we fit GLM models using either all amygdala or hippocampal category neurons recorded in a given session and compared the t-values between the two areas (only including sessions that contained category cells in both regions). This comparison revealed that the subsequent memory effect was significantly larger in the amygdala than in the hippocampus (Fig. 3b, Wilcoxon signed-rank test, *p* = 0.021). There was no significant difference between the two areas for trials in the non-preferred categories (Fig. 3d, Wilcoxon signed-rank test, *p* > 0.9), again confirming specificity to the preferred category of the cells.

Whereas the amygdala showed an early effect beginning at 350 msec, the hippocampus began to show this difference much later around 1300 msec following stimulus onset (Fig. 3e, cluster-based permutation test, *p* = 0.009). Thus, in contrast to the amygdala, the sensory and memory-predictive response did not co-occur in time in the hippocampus (in which the differential latency at which category cells first differentiate between their preferred and non-preferred category is approximately 300 msec following stimulus onset, similar to the amygdala^31,32^).

### Memory effects in the amygdala are independent of emotional valence and arousal

Previous studies have highlighted the role of the amygdala in emotional processing^34,35^, with stronger emotional content generally resulting in stronger memories. We next determine whether the emotional valence or arousal caused by the novel stimuli, rather than memory encoding per-se, gave rise to the differences in activity we observed. To do so, we gathered valence and arousal ratings for each of our images on a continuous scale ranging from −1 to 1 (Supplementary Fig. 2, see methods). We then included these valence and arousal ratings as additional fixed effects in our mixed-effects GLM model alongside memory encoding outcome (remembered vs. forgotten), with neuron ID again nested within session ID as random effects. Only preferred trials of category cells were included. This analysis revealed that the effect of memory encoding outcome on firing rate remained significant after accounting for both emotional factors (β_memory = 23.34, t = 4.51, *p* = 6.5 × 10⁻⁶). Indeed, the emotional factors (arousal and valence) did not significantly predict firing rate of category cells by themselves (β_arousal = 19.57, t = 1.18, *p* = 0.24; β_valence = −26.41, t = −1.78, *p* = 0.075). Furthermore, the relationship between firing rate and memory outcome was also present when considering the positively valenced and negatively valenced images separately (Positive valence: β = 22.29, t = 3.91, *p* = 9.2 × 10⁻^5^, Negative valence: β = 28.20, t = 2.24, *p* = 0.025). These findings suggest that the observed differences in firing rate were not due to emotional valence or arousal.

### Amygdala neural activity tracks memory strength

What aspect of episodic memory does the activity of cells in the amygdala predict? To answer this question, we compared the firing rates during encoding between two types of later remembered trials: those remembered with low and high confidence (Fig. 4a-b, example amygdala category neurons). In Fig. 4a, an example neuron is shown that shows a preference for the ‘landscapes’ category and which fired more strongly for high-confidence than low-confidence remembered images in that category. The neuron in Fig. 4b showed a similar response pattern. Population-level analysis revealed that the average firing rates of category cells in the amygdala following viewing of images belonging to the preferred category were higher for images later remembered with high confidence compared to those later remembered with low confidence (Fig. 4c, Wilcoxon signed-rank test, *p* = 3.2 × 10^−4^ for amygdala after Bonferroni correction). Time-resolved analysis revealed a graded representation of memory strength, with the highest firing rates for items remembered with high confidence, followed by low confidence, and then forgotten items. The difference in amygdala neural activity between high- and low-confidence trials was significant from 450 to 1500 msec after stimulus onset (Fig. 4d, cluster-based permutation test, *p* < 0.001). In contrast, the hippocampus showed a smaller and delayed effect (*p* = 0.012), while no significant differences observed in the other two brain regions.

**Fig. 4:**
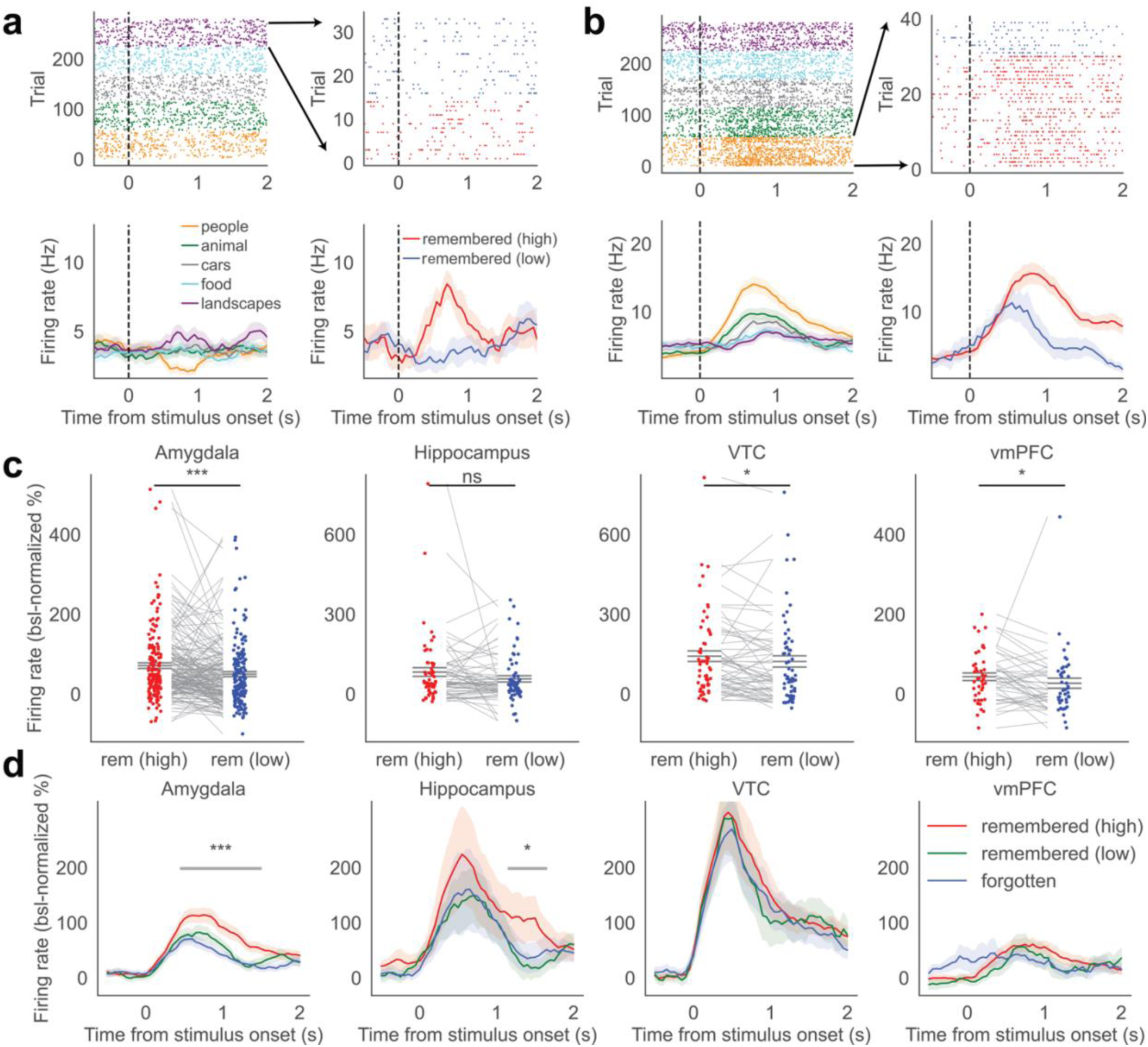
Relationship between encoding activity and memory strength. **(a-b)** Example neurons in the amygdala exhibited increased firing rates for their preferred category. Within the preferred category, neurons showed higher firing rates for high-confidence remembered items compared to low-confidence ones. **(c)** Comparison of average firing rates during the encoding period between trials that were later remembered with high versus low confidence. Each line connects the two data points from the same neuron. Statistical analysis was performed using the Wilcoxon signed-rank test. Amygdala: n = 169 neurons, *p* = 3.2 × 10^−4^; Hippocampus: n = 61 neurons, *p* = 0.59; VTC: n = 60 neurons, *p* = 0.026; vmPFC: n = 43 neurons, *p* = 0.030, *p* values were corrected for multiple comparisons. **(d)** Time-resolved firing rates for high-confidence remembered trials, low-confidence remembered trials, and forgotten trials. Statistical analysis was performed using a cluster-based permutation test. Data are presented as mean ± SEM. \**p* < 0.05, \*\**p* < 0.01, \*\*\**p* < 0.001, ns indicates non-significant.

To further assess whether firing rates during encoding predict memory strength, we used 6-level confidence ratings from the recognition part (see Fig. 1b). Ratings of 1–3 indicate decreasing certainty that the picture is new, while ratings of 4–6 reflect increasing certainty that it is old. Thus, the higher the confidence rating, the stronger the underlying memory trace. Firing rates of category-selective amygdala neurons (preferred trials only) were modeled using a generalized linear model, with memory strength (confidence ratings 1–6) as the predictor and neurons nested within recording sessions as random effects (all trials of the preferred category were included, regardless of whether they were remembered or forgotten). This analysis revealed a significant positive relationship between memory strength as indicated by confidence and firing rate during encoding (β = 6.38, t = 5.26, *p* = 1.4 × 10^−7^).

To determine whether the relationship between memory strength and firing rate in the amygdala was specific to encoding, we repeated the analysis using neural activity from the recognition part (only considering familiar stimuli, comparing those remembered and forgotten). Unlike during encoding, the firing rates of category cells (preferred trials) did not differ significantly between remembered and forgotten trials during recognition (β = 1.35, t = 0.21, *p* = 0.84), nor was there a significant relationship between firing rate and confidence ratings (during preferred trials, β = 0.95, t = 0.60, *p* = 0.55). These results suggest that the observed change in firing rate of category cells associated with memory strength is driven by processes occurring during encoding rather than during retrieval.

Our findings show that a higher firing rate of category neurons is not only associated with successful memory encoding but also with greater confidence during retrieval. This pattern suggests that increased amygdala activity supports the formation of detailed episodic memories, as opposed to familiarity-based recognition alone.

### Replication of subsequent memory effect in the classic recognition memory task

To assess the robustness of our finding, we next repeated the same analysis in a second independent dataset with a different task. Task 2 is a standard recognition memory task with two parts: a learning part and a recognition part (Fig. 5a). During the learning part, participants were presented with 100 unique, novel images (Fig. 5b). After an interval of 10-30 minutes, participants completed the recognition part, during which they viewed 100 images, with half being old (previously presented during the learning part) and the other half being new. For each image, they indicate whether it was old or new using a 6-point confidence scale (Fig. 5c). We analyzed a total of 5744 neurons (of these, 960 were category-selective) across 131 sessions from 95 subjects. Confirming our main finding, category-selective cells in the amygdala also exhibited significantly higher firing rates for remembered items compared to forgotten ones during encoding (Fig. 5d-e). Similarly, repeating our generalized linear model also showed a significant effect of memory accuracy (β = 28.35, t = 3.78, *p* = 1.6 × 10^−4^) on firing rates of category cells during preferred trials and firing rates were greater for images remembered with high confidence than for those remembered with low confidence (Fig. 5f-g). Higher confidence ratings were associated with increased neuronal firing rates (generalized linear model, β = 7.19, t = 3.96, *p* = 7.4 × 10^−5^). In contrast to the amygdala, there was no significant relationship between later remembered and forgotten trials in the hippocampus (β = 8.70, t = 1.28, *p* = 0.20 for remembered v.s. forgotten, β = 2.56, t = 1.52, *p* = 0.13 for confidence ratings). We also confirmed our findings related to emotional modulation by adding valence and arousal ratings as fixed effects to the mixed-effects GLM, along with memory encoding outcome (remembered vs. forgotten). Memory encoding outcome continued to have a significant relationship to the firing rate of amygdala category neurons after controlling for both emotional factors (β_memory = 27.31, t = 3.64, *p* = 2.7 × 10⁻^4^). Among the emotional factors, arousal but not valence significantly predicted firing rate in category cells (β_arousal = 77.65, t = 3.25, *p* = 1.2 × 10⁻^3^; β_valence = −2.27, t = −0.14, *p* = 0.89). Lastly, the association between firing rate and memory outcome was also present when analyzing positively and negatively valenced images separately (Positive: β = 22.14, t = 2.40, *p* = 0.016, Negative: β = 45.40, t = 3.66, *p* = 2.5 × 10^−4^). Thus, although arousal predicted variability in firing rates, the relationship to memory encoding outcome remained significant. This independent replication in an independent large dataset adds converging evidence to our conclusion that the activity of category-selective neurons in the amygdala is predictive of memory encoding success and underscores the generalizability of our findings across both tasks.

**Fig. 5:**
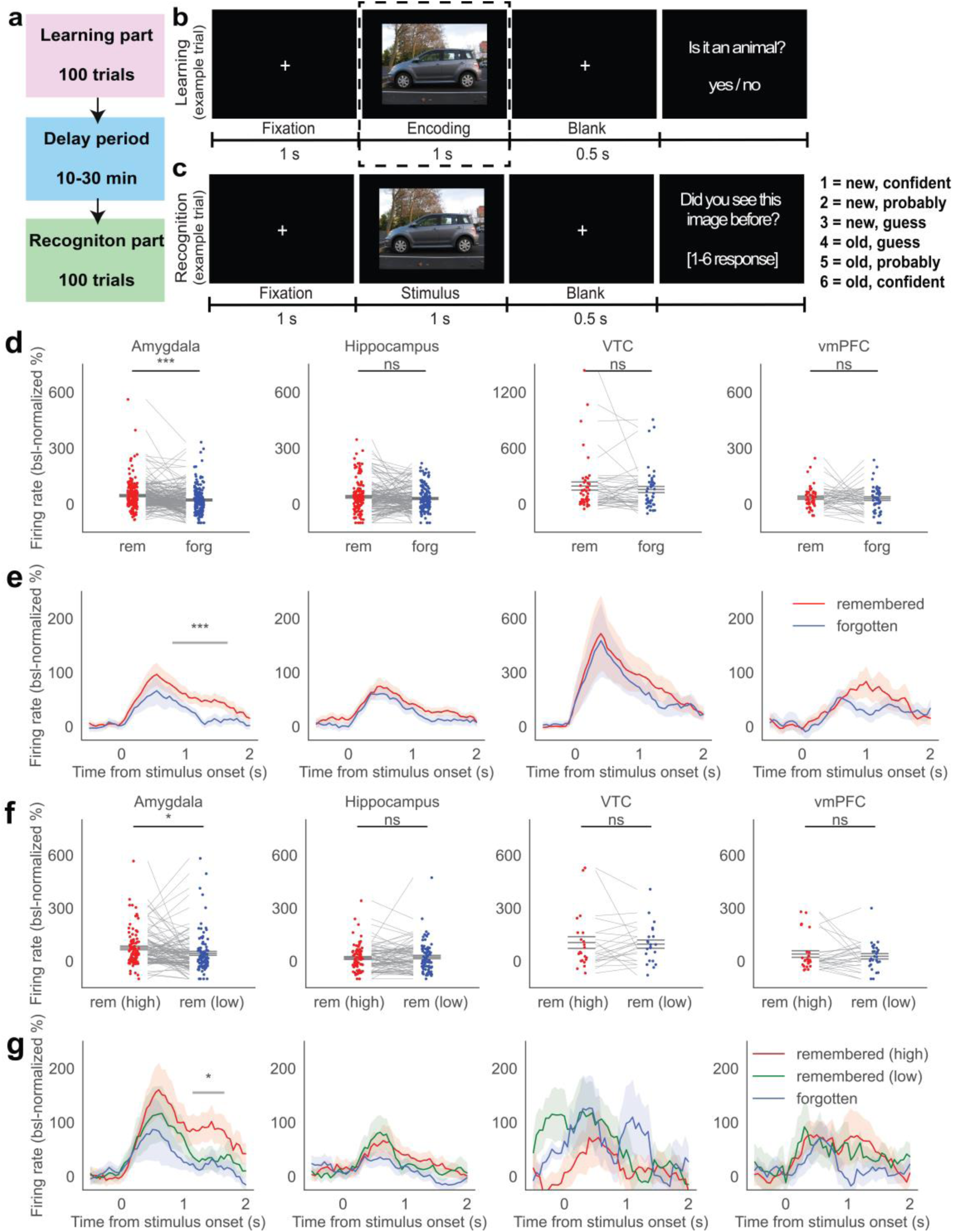
Replication of key findings in a recognition memory task (Task 2). **(a)** Task. Participants first completed a learning part, during which they viewed 100 unique images. Following a 10–30 minute delay, they proceeded to the recognition part, where they saw 100 images, with half previously seen and half new. **(b)** Learning part: On each trial, participants reported whether an animal was present in the image. **(c)** Recognition part: For each image, participants reported whether it was old or new while rating their confidence on a 6-point scale. **(d)** Comparison of average firing rates during the encoding period between trials that were later remembered versus those that were forgotten. Each line connects the two points from the same neuron. Average firing rates were significantly higher in remembered trials than forgotten trials in the amygdala (Wilcoxon signed-rank test, n = 183 neurons, *p* = 1.3 × 10^−4^ after corrected for four brain regions) but not in other regions (*p* > 0.9). **(e)** Time-resolved firing rates for remembered and forgotten trials. Statistical analysis using a cluster-based permutation test revealed a significant cluster in the amygdala (*p* < 0.001). **(f)** Comparison of average firing rates between trials later remembered with high versus low confidence. The difference was significant in the amygdala (n = 104 neurons, *p* = 0.021 after corrected for four brain regions) but not in other regions (*p* > 0.9). **(g)** Time-resolved firing rates for high-confidence remembered trials, low-confidence remembered trials, and forgotten trials. A significant cluster was identified in the amygdala (*p* = 0.014), while no significant clusters were found in other regions. \**p* < 0.05, \*\**p* < 0.01, \*\*\**p* < 0.001, ns indicates non-significant.

### Stronger subsequent memory effect in the right amygdala

Given the independent replication of our principal effect in both Task 1 and 2, we next combined both together to establish sufficient statistical power to assess potential sex and hemisphere differences in the subsequent memory effect. To do so, we fit the same mixed-effect GLM as described above (predicting firing rate as a function of whether an image was later remembered or forgotten) to all amygdala category neurons from different groups and compared the effect sizes. For sex, generalized linear models were fit separately for amygdala category neurons from female and male participants, both of which showed a significant memory effect independently (female: β = 25.41, t = 4.23, *p* =2.3 × 10^−5^, n = 203 cells, male: β = 27.85, t = 3.86, *p* = 1.1 × 10^−4^, n=156 cells). To evaluate whether effect strength differed between the two groups, we first computed the difference in t-values after randomly selecting an equal number of neurons from each group (based on the smaller group size). The observed difference was obtained by averaging across 1000 subsampling iterations. To evaluate significance, sex labels were randomly permuted 1000 times to generate a null distribution, and the observed difference was then compared against this distribution. No significant difference between sexes was observed (Supplementary Fig. 3a). For hemispheric comparisons, the same procedure was applied to amygdala neurons from the left and right hemispheres. The subsequent memory effect was significant in both hemispheres (left: β = 19.18, t = 2.64, *p* = 8.2 × 10^−3^, n = 183 cells, right: β = 30.44, t = 5.56, *p* = 2.7 × 10^−8^, n = 198 cells), indicating that the firing rate of category neurons in both the left and right hemispheres predicted the success of memory encoding. We compared the effect sizes between the two hemispheres with the same procedure as described for sex differences. This revealed that the effect was significantly stronger in the right compared to the left hemisphere (*p* = 0.009, Supplementary Fig. 3b-c). This suggests that amygdala neurons in the right hemisphere contribute more strongly to the encoding of the types of memories (visual images) examined in our tasks.

### Phase locking is stronger during viewing of later remembered items

Why does increased firing of category cells lead to stronger memories? One possibility is that if firing is stronger, coordination of spike timing with ongoing local field potential oscillations is stronger due to higher levels of overall synaptic activity. This hypothesis is motivated by the finding that strong spike-field coordination can lead to stronger plasticity^36^ and thus memory, in particular if coordination is between spiking and field potential activity in different brain areas^37–39^. To test this hypothesis, we next tested the extent of spike-field coherence of category cells. We began by investigating whether the spike timing relationship between local field potentials and spikes of category cells within the amygdala predicted successful memory encoding in Task 1 during the encoding period (Fig. 6a; this analysis was only possible in Task 1 due to lower trial numbers in Task 2). We found that spike-field coherence (SFC) within the amygdala was significantly stronger during trials in which items were later remembered compared to forgotten trials, but again only for preferred-category trials of category cells (Fig. 6b, cluster-based permutation test, 9-18 Hz, *p* < 0.001; see below for statistics for non-preferred trials). In contrast, no significant differences were found in the hippocampus (Fig. 6c).

**Fig. 6:**
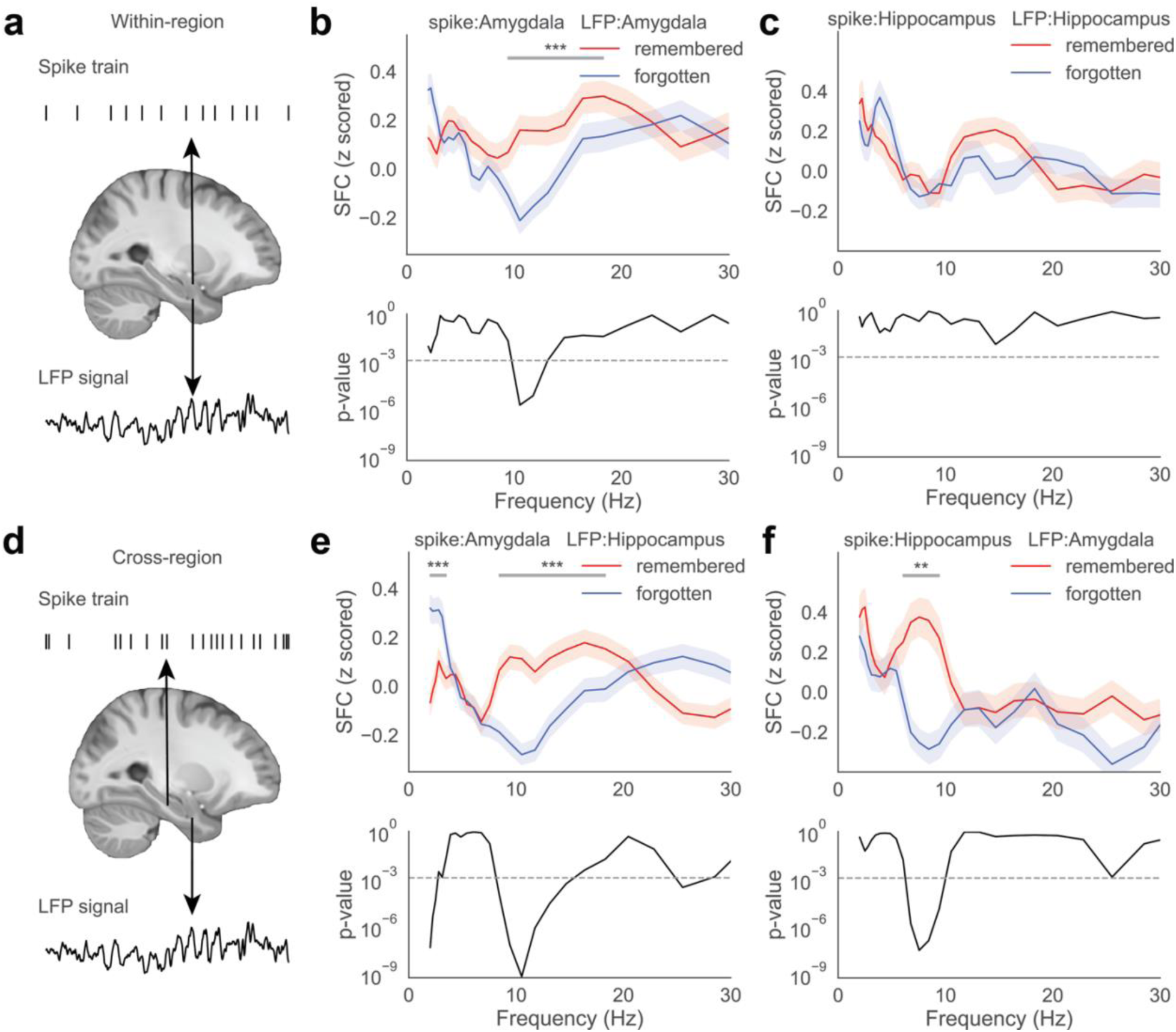
SFC differentiates between trials that were subsequently remembered and those that were forgotten. **(a)** A schematic of spike-field coherence between spikes and LFPs within the same brain region. **(b)** SFC between spikes and LFP signal in the amygdala. The upper plot shows z-scored SFC across frequencies, presented as mean ± SEM. Statistical analysis was performed using a cluster-based permutation test. Category neurons in the amygdala were more closely aligned with local LFP during later remembered than forgotten trials of the preferred category (n = 346 combinations for n = 48 category cells, *p* < 0.001). The lower plot displays *p*-values obtained from the GLM model, with the peak difference between remembered and forgotten trials (corresponding to the largest t-value and smallest *p*-value) observed at 10.5 Hz. **(c)** SFC between hippocampal category neurons and LFPs did not differ significantly between later remembered and forgotten trials (n = 203 combinations). **(d)** A schematic of long-range SFC between spiking activity and LFPs across different brain regions. **(e)** Spike timing of category neurons in the amygdala was more synchronized with hippocampal activity during successful encoding, with the peak difference between remembered and forgotten items at approximately 10.5 Hz (n = 460 combinations, *p* < 0.001). **(f)** Spike timing of category neurons in the hippocampus was more synchronized with amygdala activity in remembered trials, showing a peak difference at 7.6 Hz (n = 141 combinations, *p* = 0.002).

Next, we tested whether the extent of inter-areal coordination between the amygdala and hippocampus during encoding was predictive of encoding success. Across the amygdala and hippocampus, category-selective neurons demonstrated greater phase locking to ongoing low-frequency activity during successful compared to unsuccessful memory encoding (Fig. 6d-f). In the amygdala, the spike timing of category neurons aligned more closely with hippocampal LFP activity during remembered trials (Fig. 6e, 8-18 Hz, *p* < 0.001). Similarly, spike timing of category neurons in the hippocampus was more synchronized with LFP activity in the amygdala during successful encoding (Fig. 6f, 6-9 Hz, *p* = 0.002). These findings suggest that low-frequency oscillatory activity not only structure local spiking activity within the amygdala, but in addition also coordinate neural activity across the amygdala-hippocampal circuit to support memory formation.

Lastly, we repeated the same analysis using trials from the non-preferred category. While a weaker memory effect within the amygdala remained, no significant coordination (SFC) between the amygdala and hippocampus was seen during the encoding of non-preferred items (Supplementary Fig. 4). Similarly, when analyzing all trials regardless of category preference, no significant coordination was observed. These findings suggest that the effect is specific to trials in which the neuron is actively engaged in processing the stimulus, underscoring the functional specificity of these interactions.

## Discussion

Among the vast amounts of new information we experience and process every day, we remember only a very small subset. What determines whether an event, at the moment it happens, is encoded into long-term memory? While the hippocampus is essential for episodic memory formation^24,40^, the extent to which hippocampal encoding processes are engaged is modulated extensively by the interplay of the hippocampus with other cortical and sub-cortical brain areas. Here, we revealed one such modulatory process: category-selective neurons in the amygdala, which fire earlier and more strongly for items that are later remembered compared to those that are later forgotten. This early predictive signal in the amygdala reveals a cellular-level understanding for how the amygdala might initiate successful memory encoding for a broad class of visual stimuli. In contrast, prior research has emphasized the modulatory role of the amygdala in enhancing hippocampal-dependent memory only for emotionally highly salient and charged stimuli^25,38^. For example, imaging research has shown that individuals with greater amygdala activity during encoding have better memory for emotional than for neutral stimuli, while lesion evidence indicates that bilateral amygdala damage impairs declarative memory for emotional material^13,41^. Instead, our findings indicate that the amygdala plays a much broader role in memory formation extending much beyond its traditional role in processing emotional salience.

Our study provides novel evidence for the amygdala’s role in rapid single-trial learning from single exposures to complex visual stimuli, a type of learning that humans excel at^42^. While humans excel in such learning, the coding principles and neural mechanisms that enable such fast plasticity without causing any apparent catastrophic interference remain largely unknown^43–45^. Previous research has shown that concept cells in the amygdala exhibit higher firing rates when locations of familiar items from the preferred concept of the cell are accurately remembered after a brief 15-second distraction in an associative memory task, suggesting a role in encoding spatial information in short-term memory^46^. In contrast to this paradigm, our study explores how the amygdala modulates the encoding of novel stimuli into long-term memory (i.e. a prolonged delay). We found that the activity of category-selective neurons in the human amygdala during encoding reliably predicts subsequent long-term memory performance. These neurons responded more strongly to items that were later remembered, but only when the items belonged to their preferred category. The specificity of this effect to images belonging to the preferred category of the cells indicates that the amygdala was involved in engaging stimulus-specific circuits for memory formation, rather than more general arousal processes that would be expected to affect all stimuli rather than a specific subset. Prior work has reported category-general memory signals, such as increased high-frequency activity for emotional words and shared responses across stimulus types like faces and scenes during successful memory encoding^25,47^. In line with these findings, we observed a subsequent memory effect when considering the mean firing rates of all recorded amygdala neurons following stimulus onset. However, this effect was much weaker than when only considering category cells, indicating that it might be masking a much stronger effect that is present only in category cells, as opposed to a nonspecific population-level effect that may arise from domain-general attention or arousal differences. Confirming the former interpretation, when we repeated our analysis including all cells but excluding for each cell the trials belonging to the category for which it fired most, the effect became non-significant. This analysis indicates that the mean firing rate differences were primarily due to responses of category cells to their preferred category. Thus, our findings reveal a stimulus-specific component of memory encoding in the amygdala.

We found that amygdala activity during encoding was greater for items later remembered with high confidence compared to those remembered with low confidence. According to signal detection theory, recognition decisions are based on evaluating continuous memory signals, with confidence ratings likely reflecting the strength of these signals^48,49^. Consistent with this theory, previous work has shown that memory-selective neurons, which distinguish between novel and familiar stimuli during retrieval, exhibit activity that correlates with confidence during retrieval but not during encoding^31^. Our current results demonstrate that during memory encoding, a different group of neurons in the amygdala, which are category-selective, track the strength of the resulting memory traces of the ongoing memory encoding process. The activity of these neurons during encoding predicts the confidence during later retrieval of the same stimulus, suggesting a role of category cells in the amygdala in the formation of episodic memories beyond familiarity-based recognition^50–52^.

Although interactions between the amygdala and hippocampus have long been associated with memory encoding processes^38,39,53^, how single neurons synchronize with LFPs across these brain areas during memory encoding remains elusive^38,54–57^. We observed that phase locking of spikes fired by category cells with LFP in the amygdala and hippocampus was significantly stronger during the encoding of subsequently remembered items. This effect was again specific to only the images that belonged to the preferred category of a cell, further highlighting the specificity of the circuit engaged by specific images. This enhanced phase locking implies that these regions communicate more effectively when information is successfully being encoded. Previous studies reported that stronger within-region spike field coupling in the amygdala and hippocampus predicts memory encoding success for images and words^36,58^. In contrast, a spatial memory study found robust theta phase locking of neurons but no difference in SFC between later remembered and forgotten locations, which may have been because the task tested spatial memory and/or the short interval between encoding and retrieval^59^. Here we found a subsequent memory effect in within-region SFC in the amygdala, but not in the hippocampus, suggesting that distinct medial temporal regions may differentially contribute to encoding. Prior studies either pooled neurons between the two areas^36,59^ or excluded the amygdala entirely^58^, so did not identify that amygdala (but not hippocampal) activity is strongly predictive of encoding success. Beyond local SFC, we also observed that interregional SFC between the amygdala and hippocampus predicts memory encoding success, thereby extending prior evidence of within-region phase locking in the medial temporal lobe. Temporal coordination between amygdala rhythms and hippocampal spiking may help optimize synaptic plasticity through mechanisms such as spike-timing-dependent plasticity^60–62^. Overall, these results underscore the critical role of low-frequency neural activity in temporally organizing distributed neuronal processes to support effective memory encoding. The stimulus specificity of the observed phase locking, as opposed to a more general effect, supports the idea that episodic memories depend on the coordinated activation of assemblies of memory-content selective cells^63,64^. This synchrony may facilitate the rapid formation of associative links between these assemblies, providing a mechanism for binding related representations into coherent episodic memory traces^65,66^.

In summary, our findings support the hypothesis that the amygdala plays a broad role in memory encoding, extending much beyond its traditionally recognized function in emotion and social behavior^28,67,68^. Our findings are compatible with the well-recognized effects of amygdala impairment in memory disorders: the amygdala is affected early in the progression of Alzheimer’s disease, and the magnitude of its atrophy has been associated with overall disease severity^69,70^. Also, several studies have demonstrated that amygdala volume predicts memory performance, even after accounting for other medial temporal lobe structural measures^71,72^. Going beyond structural associations, our finding that amygdala activity during encoding predicts subsequent memory performance underscores that the amygdala is a critical contributor to episodic memory formation. This has important clinical implications, as the memory impairments seen in disorders such as Alzheimer’s disease and temporal lobe epilepsy might be partly due to disrupted encoding processes within the amygdala. Therefore, our findings indicate that the amygdala might be a suitable therapeutic target for treating memory disorders, for example through deep brain stimulation^14,15,73,74^ or functional ultrasound stimulation^75,76^. Such an approach is of particular interest given that direct stimulation of the hippocampus has frequently been reported to impair, rather than enhance, memory performance^20,21,77^. Our findings also highlight a major obstacle that has to be overcome for this approach to become effective: stimulation has to be selective to the cells which are excited by a given stimulus (and not all cells in general). Recent work on electrical stimulation highlights that this might be feasible: depending on the stimulation protocol, electrical stimulation can be remarkably selective to certain cell types^78^, and our recent work shows that low-amplitude stimulation can selectively engage only excited cells^79^, which in the present case would be category cells when exposed to a stimulus belonging to their preferred category. We found that while amygdala category neurons in both hemispheres encoded memory-related information, the effect was stronger in the right amygdala. This hemispheric asymmetry may be driven by our use of visual stimuli as memoranda, as previous studies indicate that left medial temporal lobe damage (including the amygdala) impairs verbal memory, whereas right-sided damage disrupts nonverbal visual memory^80,81^. This hemispheric difference can also be seen as a function of the location of the seizure onset zone in patients^82^. Previous fMRI work has also shown that right, but not left, amygdala activity predicts the subjective vividness of memory^83^. Consistent with this, our observation that the subsequent memory effect is stronger in the right amygdala suggests that targeting the right amygdala may be the best stimulation strategy for enhancing memory, particularly for nonverbal information.

Taken together, our findings highlight the amygdala as a promising focus for future research and therapeutic strategies aimed at enhancing memory and treating memory-related disorders.

## Methods

### Participants

We studied two patient cohorts who performed different memory tasks during intracranial monitoring for drug-resistant epilepsy. All patients were implanted with depth electrodes for seizure localization as part of their clinical evaluation. Implantation sites were determined according to clinical need. All participants provided informed consent and volunteered for the study they participated in. Recordings were conducted at Cedars-Sinai Medical Center, Toronto Western Hospital, and Johns Hopkins Hospital. Studies received approval from the Institutional Review Board of the respective institution where each patient was enrolled. The Task 1 cohort includes 42 patients (21 females, 20 males, and 1 non-binary, age 39.5 ± 12.8 years). A subset of data from the medial temporal region of Task 1 has been previously published^84^. The Task 2 cohort includes 95 patients (51 females, 43 males, 1 non-binary; age 37.2 ± 14.0 years). For task 2, 59 patients were part of a previously published dataset^85^, and the remaining 36 patients represent newly collected data not published so far.

### Tasks

Task 1 was a working memory task followed by a recognition memory test (Fig. 1a). The working memory part of this task included 140 trials, each starting with a fixation cross shown for 0.9 to 1.2 seconds. Depending on the memory load condition, this was followed by either a single image or a sequence of three images, with each image appearing on screen for 2 seconds. All images shown during this encoding period were novel to the patient. After the image presentation, there was a 2.55 to 2.85 second maintenance period, with the word “HOLD” displayed on the screen. A probe image was then presented, and participants used a Cedrus response pad to indicate whether it matched any image from the current trial. For patients participating in multiple sessions, a completely new set of images was introduced in each session to guarantee the novelty of all images across sessions. These images belonged to one of five categories: people, food, animals, cars, or landscapes, and were presented at a visual angle of 10.5° × 7°. Patients then completed the long-term memory part of the task after a delay of 10 to 30 minutes following the completion of the working memory part (Fig. 1a). During this part, 400 images were presented one by one, with 200 being old (previously shown in the working memory part) and 200 being new (never shown so far). The new images were chosen from the same five visual categories as the old images. Each trial started with a fixation cross displayed for 0.9 to 1.2 seconds, followed by an image. Patients were asked to indicate both their choice and confidence in the decision simultaneously by selecting a number from 1 to 6 on a six-button response pad. Buttons 1–3 were assigned to ‘new’ responses and buttons 4–6 were assigned to ‘old’ responses with increasing confidence. The image remained on the screen until a response was made.

Task 2 was a classic ‘new/old’ recognition memory task. This task consisted of two parts: a learning part and a recognition part, separated by a 10–30 minute delay (Fig. 5a)^31^. During the learning part, 100 unique and novel pictures were shown. To ensure participants remained engaged and attentive, they were asked to determine whether an animal was present in the picture following each learning trial. During the recognition part, participants were shown 100 pictures: 50 that had appeared previously in the learning part (old) and 50 novel ones they had not seen before (new). As in the other task, participants were asked to indicate whether it was ‘old’ or ‘new’ and to select their confidence level for each picture by choosing a number from 1 to 6. Performance feedback was provided only after the experiment was completed. Pictures from five different categories were presented at a visual angle of 9° × 9°. An equal number of pictures from each category was displayed during the recognition part to mitigate bias in memory towards certain categories.

### Behavioral analysis

Receiver operating characteristic (ROC) curves were constructed by calculating the true positive rate (the proportion of old items correctly identified as old) and the false positive rate (the proportion of new items incorrectly identified as old) at each confidence threshold. These values were used to plot curves showing the trade-off between hit rates and false alarm rates across confidence levels, with the false positive rate on the x-axis and the true positive rate on the y-axis. The area under the curve (AUC) was computed as a summary measure of discriminative ability across all possible thresholds, ranging from 0.5 (chance performance) to 1.0 (perfect classification), with higher values indicating better memory performance. Data from two sessions were excluded from all analyses due to performance falling below 55% accuracy in either the working memory or long-term memory part.

### Electrophysiology

We recorded neural activity using hybrid depth electrodes, each contained eight microwires at the tip of the electrode shank. Broadband signals were recorded at 32 kHz (ATLAS system, Neuralynx; Cedars-Sinai Medical Center and Toronto Western Hospital) or 30 kHz (Blackrock Neurotech; Johns Hopkins Hospital) from each microwire. Each recording site used a local reference, selected from either one of the eight microwires or a dedicated low-impedance reference channel included in the bundle if all other channels captured neuronal spiking activity.

### Electrode localization

Electrode locations were determined by co-registering pre-operative magnetic resonance imaging (MRI) with post-operative MRI or computed tomography scans using Freesurfer^86^. The anatomical location of these electrodes was verified by visually inspecting patient-specific imaging. We then aligned each patient’s pre-operative MRI to the CIT168 template brain in MNI152 coordinates as previously described^87–89^. The MNI coordinates of electrodes across all patients were plotted onto this template brain.

### Spike detection and sorting

The raw signal was filtered offline with a zero-phase filter in the 300-3000 Hz band. Spikes were detected and sorted using the semiautomated template-matching algorithm Osort v4^90^. We assessed the quality of spike sorting for all putative single units using standard metrics, such as the signal-to-noise ratio of the waveform, the proportion of inter-spike intervals shorter than 3 msec, and the projection distance between isolated cluster pairs^91^. Only well-isolated neurons as determined by these metrics were included in the subsequent analysis.

### Selection of category neuron

To identify neurons that exhibited systematic differences in their responses to distinct picture categories, we calculated the firing rate within a 200 to 1000 msec window following stimulus onset in each trial (all encoding periods and the probe period). Firing rates were then grouped according to the category of the presented picture. To assess category selectivity, we conducted a 1×5 ANOVA for each neuron, using picture category as the grouping variable. This was followed by a one-sided permutation-based t-test, comparing the category with the highest firing rate to all other categories (1000 permutations). A neuron was classified as category-selective if both statistical tests reached significance (*p* < 0.05). We refer to the category with the highest average firing rate as the neuron’s preferred category.

To evaluate whether the number of category-selective neurons identified in each brain area was larger than expected by chance, we repeated this selection process 1000 times after randomly shuffling category labels across trials. For each iteration, we obtained the proportion of category-selective neurons relative to the total number of neurons within each region. These 1000 values formed the empirically derived null distribution for the proportion of category neurons that would be expected by chance. We considered the number of category-selective neurons observed in a given brain region significant if the actual proportion was higher than 99% of the null distribution (*p* < 0.01).

### Generalized linear mixed-effects model

We used a generalized linear mixed-effects model to assess whether firing rate predicts memory encoding success. In our model, memory outcome (remembered, forgotten) was treated as a fixed effect, while neuron ID nested within session ID were included as random effects. The model was specified as: Firing rate ∼ memory outcome + (1 | session/neuron). To test whether firing rate during the encoding period predicts memory strength, we used confidence rating (6 levels, from 1 to 6) in the later recognition part as memory strength for each item. Ratings of 1, 2, and 3 indicate that the patient is sure, less sure, or very unsure that the picture is new, whereas ratings of 4, 5, and 6 indicate increasing certainty that the picture is old. Therefore, higher confidence ratings are taken to reflect a stronger memory. The model used for this analysis was: Firing rate ∼ confidence rating + (1 | session/neuron). Firing rate during the encoding period was normalized as a percentage change relative to the baseline period (−0.9 to −0.3 seconds before the onset of the first picture). Neurons with a baseline firing rate below 0.2 Hz were excluded to prevent normalization artifacts. The *p*-value for each parameter was obtained using a two-sided statistical test.

### Emotional rating collection and analysis

To determine whether differences in firing rate are, at least partially, driven by emotional responses evoked by the images rather than memory encoding success, we collected valence and arousal ratings for all images from an independent sample of subjects (online, supplementary Fig. 2a). The experiment was implemented using PsychoPy, hosted on Pavlovia (https://pavlovia.org). 40 participants were recruited through Prolific (https://www.prolific.com, 15 females, 24 males, and 1 unknown, age 46.6 ± 15.0 years). All individuals provided informed consent prior to participation and received monetary compensation. They were asked to indicate for each image how viewing it made them feel. They provided their response by clicking on a scale bar ranging from −1 to 1. Subjects provided ratings for valence (how pleasant you feel while looking at it, from very unpleasant to very pleasant) and arousal (how aroused you feel while looking at it, from calm/relaxed/unaroused to excited/wide-awake/aroused). To ensure rating reliability, 20 images were repeated for each participant. We included data only from the 24 participants who showed reliable ratings, defined as a significant Pearson correlation between the two presentations of the same image for both valence and arousal ratings (average correlation for valence = 0.84, average correlation for arousal = 0.75). The average valence and arousal ratings for each image were then used as additional predictors in the respective mixed-effects models alongside the memory outcome (remembered vs. forgotten) predictor.

### Population decoding (only Task 1)

We performed population decoding using a pseudo-population constructed by combining neurons across all recording sessions. We first identified all category-selective neurons in the amygdala that had at least ten trials in their preferred category for both remembered and forgotten conditions. For each of these neurons, we randomly selected firing rates from ten trials per condition, ensuring an equal number of samples for both remembered and forgotten trials. These firing rates were then concatenated across neurons to build a pseudo-population. To assess classification performance, we used a stratified 10-fold cross-validation scheme, ensuring that each fold contained the same proportion of remembered and forgotten trials. Before classification, firing rates for each neuron were standardized across trials using the mean and standard deviation computed from the training set. Classification was then carried out using a support vector machine with a linear kernel, implemented via the SVC class from Python’s Scikit-learn library. This decoding process was repeated 1000 times to generate a stable and reliable estimate of classification accuracy. For statistical evaluation, the *p*-value was defined as the proportion of instances in which chance performance exceeded the observed classification accuracy.

### Preprocessing of local field potentials (only Task 1)

To prevent contamination of local field potentials by spiking activity, we removed spike waveforms by linearly interpolating the raw signal from −1 to 2 msec relative to each spike onset^92^ in the raw data before downsampling. The signal was then low-pass filtered at 175 Hz using a zero-phase filter and downsampled to 400 Hz. To reduce the effects of line-noise artifacts, we applied zero-phase band-stop filters at 59.5–60.5 Hz and at its first harmonic, 119.5–120.5 Hz. The processed signal was visually inspected, and trials containing artifacts were excluded. Any channels or brain regions exhibiting substantial noise or signs of epileptic activity were completely excluded from further analysis.

### Spike field coherence (only Task 1)

To assess the relationship between the spike timing of category-selective neurons and the LFP phase, we calculated spike-field coherence (SFC) across all neuron-to-channel pairs within the ipsilateral hemisphere, both within and between brain regions. To minimize filter-induced edge artifacts, we first extracted data segments from −1000 to 3000 msec relative to the stimulus onset across all clean trials and channels. Consistent with previous work, we then applied a complex Morlet wavelet convolution to extract the instantaneous phase of the LFP signals^92,93^.

This was done using a continuous wavelet transform with 40 complex Morlet wavelets, logarithmically spaced between 2 and 150 Hz. The number of cycles per wavelet also increased logarithmically with frequency, ranging from 3 to 10 cycles. We then cut the trials to an analysis window of 200 to 1000 msec following stimulus onset. To ensure a sufficient number of spikes, we included only neurons with at least twenty spikes in both remembered and forgotten trials. To prevent bias due to unequal spike counts, we randomly subsampled spikes from the larger group to match the number in the smaller one^36^. For each spike, we identified the nearest LFP phase based on its timestamp, then computed the mean vector length by averaging these phases in polar space across all spikes. This process was performed separately for all frequencies. To ensure robustness, we repeated the subsampling procedure 500 times and averaged the SFCs across all repetitions.

To normalize the SFC for each neuron-to-channel combination, we created a surrogate distribution by inducing random noise to the spike timestamps within each condition and repeated this process 500 times. This approach reduced any potential biases in the SFC that could arise from systematic differences between conditions such as the spectral power of a specific frequency. We then fitted a normal distribution to the surrogate data to obtain the mean and standard deviation, which were used to z-score the raw SFC for each condition. To compare the z-scored SFC between remembered and forgotten trials within the preferred category, we applied cluster-based permutation statistics to identify frequency ranges with significant differences. This analysis was then repeated for the non-preferred category.

## Data and Code Availability

Data and code that support the findings of this study will be made publicly available upon acceptance.

## Acknowledgments

We thank the clinical teams at Cedars-Sinai Medical Center (in particular, Chrystal Reed, Jeffrey Chung, and Lisa Bateman), Toronto Western Hospital, and Johns Hopkins School of Medicine for their support in patient care and data acquisition. We thank Ralph Adolphs for advice and the members of the Adolphs and Rutishauser laboratories for discussions. We are deeply grateful to all patients who participated in this study. This work was supported by the BRAIN Initiative through the National Institutes of Health (grant no. U01NS117839 to U.R.), and a Center for Neural Science and Medicine Postdoctoral Fellowship by Cedars-Sinai (to J.X.).

## Contributions

J.X. and U.R. conceived the project. J.X., J.D., Y.S., N.K. and C.M. collected the data. J.X. performed data analyses. W.S.A., T.A.V., and A.N.M. provided patient care and performed surgeries. J.X. and U.R. wrote the original manuscript, and all authors contributed to its review and editing. J.X. and U.R. acquired the funding.

## Competing interests

The authors declare no competing interests.

## Supplementary figures

**Supplementary Fig. 1:**
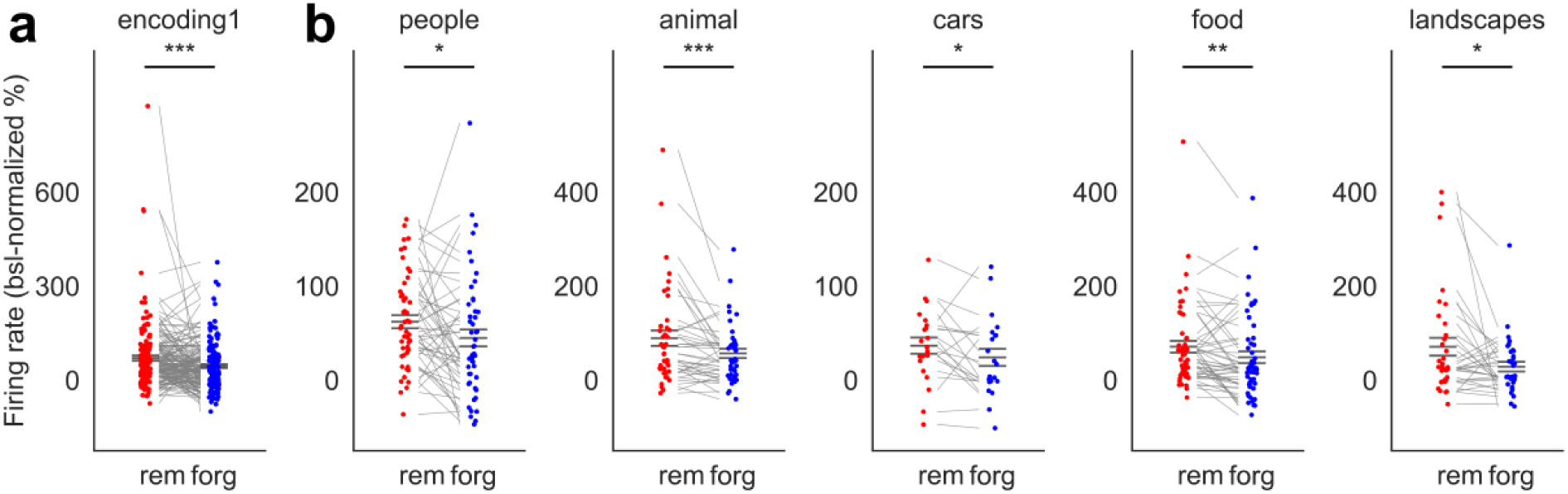
Comparison of remembered and forgotten trials in the amygdala for different subsets of trials. **(a)** Category neurons in the amygdala exhibit elevated firing rates during successful encoding of preferred-category trials, when considering only encoding1 trials (*p* < 0.001). Each line connects the two dots belonging to the same neuron. Statistical analysis was performed using the Wilcoxon signed-rank test. **(b)** The difference between remembered and forgotten trials also holds when analyzing category cells with different preferred categories separately, across all five categories (*p* < 0.05 for all categories). For each panel, only the subset of category cells with the stated preferred category was included (people: n = 50, animal: n = 41, cars: n = 21, food: n = 52, landscapes: n = 34). \**p* < 0.05, \*\**p* < 0.01, \*\*\**p* < 0.001.

**Supplementary Fig. 2:**
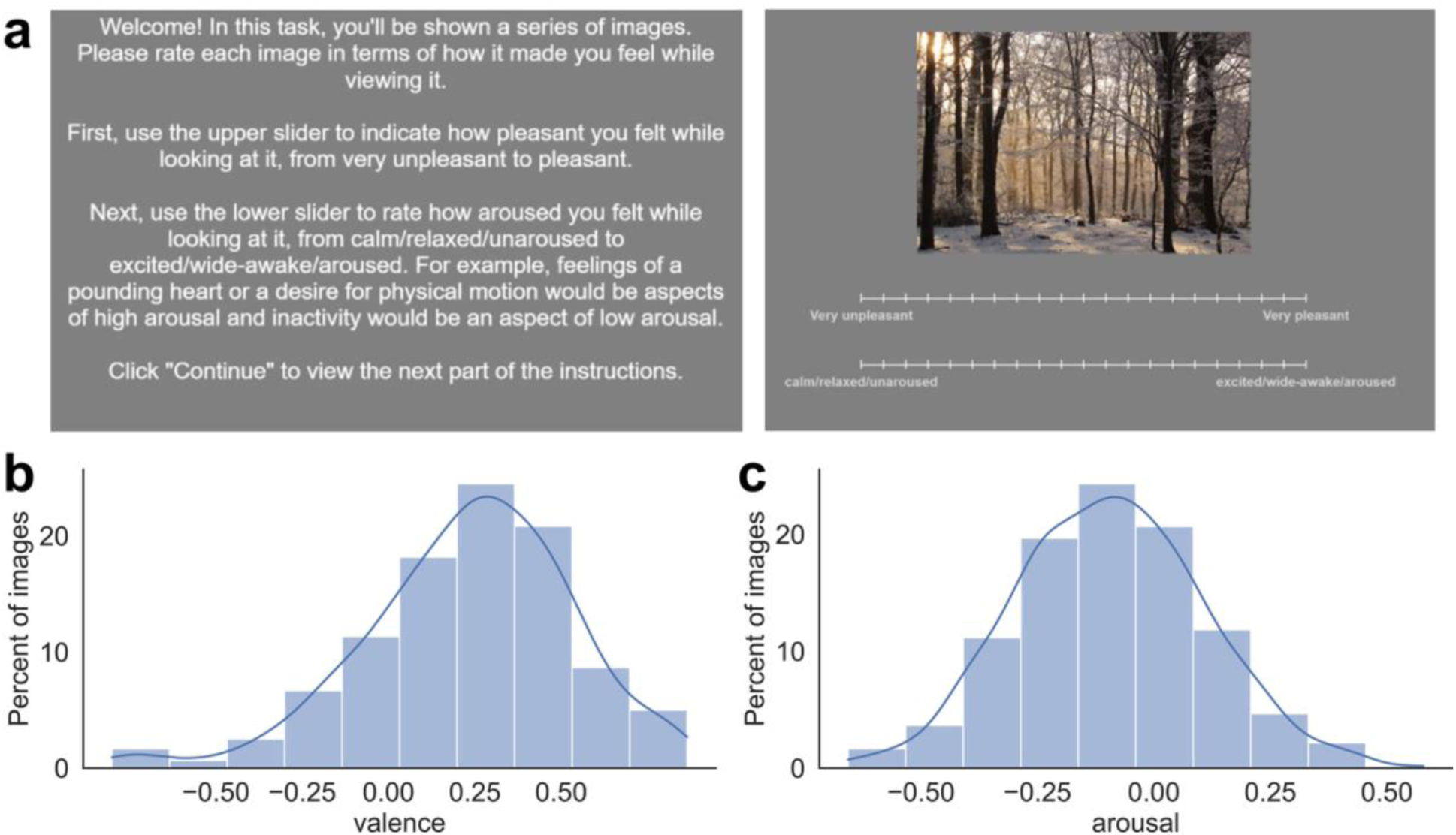
Emotion rating task. **(a)** Example screens from the task. Valence and arousal ratings were collected for all images from an independent online sample. Participants indicated how each image made them feel by clicking on a scale bar. **(b)** Distribution of valence ratings. **(c)** Distribution of arousal ratings.

**Supplementary Fig. 3:**
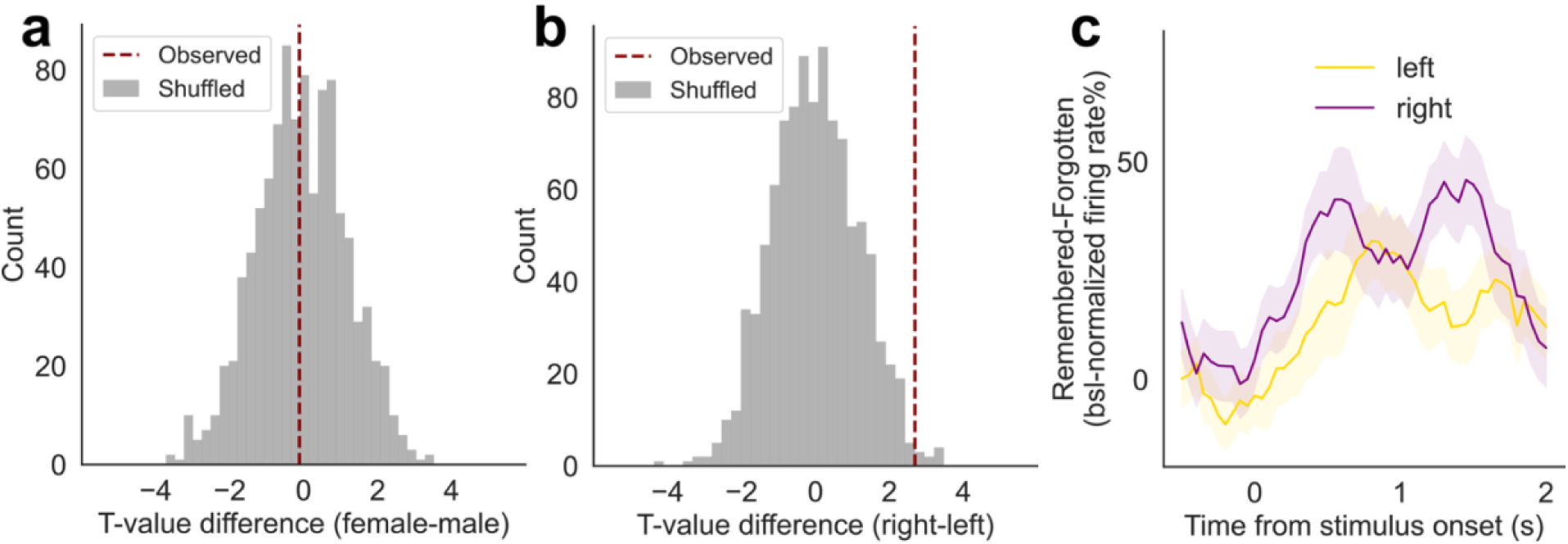
Permutation tests for sex and hemispheric differences in the subsequent memory effect. **(a)** Sex differences. Shown is the null distribution of t-value differences between female and male patients, generated from 1000 random permutations of sex labels (grey). The observed true difference (red) did not differ significantly from the null distribution. **(b)** Right vs. left hemispheric differences. Shown is the null distribution of t-value differences between amygdala category neurons from the right and left hemispheres (grey, 1000 permutations). The observed difference (red dashed line) was significantly greater than chance (*p* = 0.009), reflecting a significantly stronger effect in the right amygdala compared to the left amygdala. **(c)** Illustration of the hemispheric difference shown in panel b. Average time course of firing rate differences between subsequently remembered and forgotten images across all neurons, considering only the responses during each neuron’s preferred category. Responses are shown separately averaged across all category-selective amygdala neurons in the left and right hemispheres, confirming that responses are stronger on the right side. Data are presented as mean ± SEM.

**Supplementary Fig. 4:**
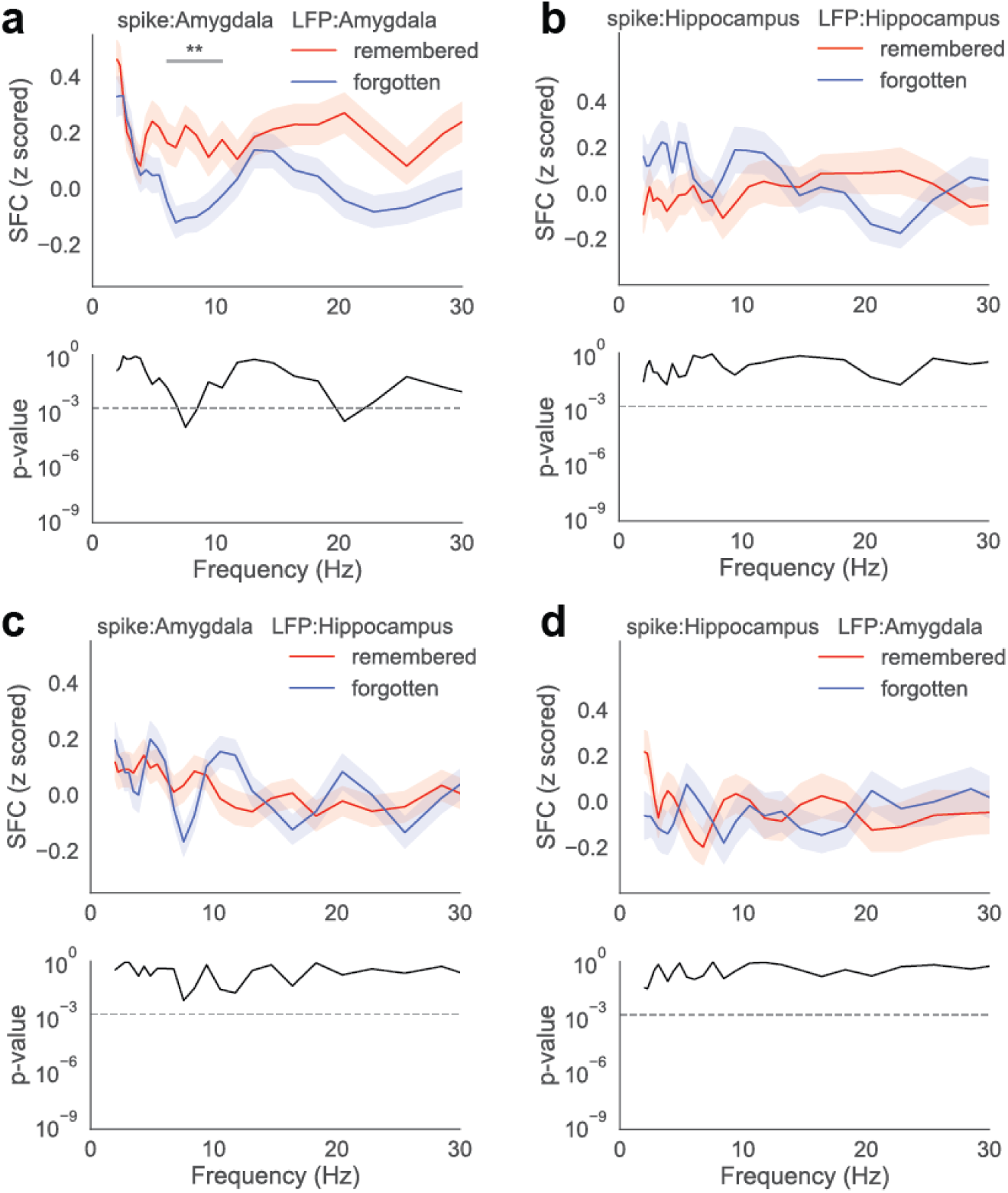
Coordination between amygdala and hippocampus is absent during encoding of items in non-preferred category. **(a)** Spike-field coherence between spikes and LFPs within the amygdala were stronger during trials that were later remembered (cluster-based permutation test, n = 279 combinations, frequency = 6–11Hz, *p* = 0.004). The upper plot shows z-scored SFC across frequencies as mean ± SEM, while the lower plot shows *p*-values derived from the GLM analysis. **(b)** There was no significant difference within the hippocampus (n = 143 combinations). **(c-d)** Cross-region SFC between category neurons and LFPs did not differ significantly between later remembered and forgotten trials (n = 334 and n = 113 combinations for (c) and (d), respectively).

## Notes

### Competing Interest Statement

The authors have declared no competing interest.

